# Duodenal organoids from metabolic dysfunction-associated steatohepatitis patients exhibit altered digestive homeostasis

**DOI:** 10.1101/2024.07.02.601648

**Authors:** Alia Hadefi, Morgane Leprovots, Gilles Dinsart, Maryam Marefati, Marjorie Vermeersch, Daniel Monteyne, David Pérez-Morga, Anne Lefort, Frédérick Libert, Laurine Verset, Claire Liefferinckx, Christophe Moreno, Jacques Devière, Eric Trépo, Marie-Isabelle Garcia

**Affiliations:** IRIBHM, Jacques E. Dumont, Faculty of Medicine, Université Libre de Bruxelles ULB, Route de Lennik 808, 1070, Brussels, Belgium; Department of Gastroenterology, Hepatopancreatology, and Digestive Oncology. CUB Hôpital Erasme, Hôpital Universitaire de Bruxelles (HUB), Université Libre de Bruxelles, Brussels, Belgium; Laboratory of Experimental Gastroenterology, Université Libre de Bruxelles, Brussels, Belgium; Center for Microscopy and Molecular Imaging, Université Libre de Bruxelles (ULB), Charleroi, Belgium; Université Libre de Bruxelles, BRIGHTcore ULB-VUB and Institute of Interdisciplinary Research in Human and Molecular Biology (IRIBHM), Brussels, Belgium; Institut Jules Bordet, Hôpital Universitaire de Bruxelles (HUB), Centre d’Anatomie pathologique, rue Meylermeersch, 90, 1070 Brussels – Belgium

**Keywords:** MASH, stem cells, cell adhesion, organoids, duodenum, intestine, permeability

## Abstract

**Background and Aims:** Metabolic dysfunction-associated steatohepatitis (MASH) is a progressive liver disease that can lead to fibrosis, cirrhosis, and hepatocellular carcinoma. Though MASH is closely tied to metabolic risk factors, the underlying pathogenic mechanisms remain scarcely understood. Recent research underscores the importance of the gut-liver axis in its pathogenesis, an aspect less explored in human studies. Here, we investigated whether the duodenal epithelium of MASH patients, could exhibit intrinsic dysfunctions.

**Methods:** Duodenal epithelial organoids were generated from 16 MASH patients and 14 healthy controls. Biopsies and patient-derived organoid transcriptomes were then analyzed to evaluate if specific intestinal pathways were differentially modulated in MASH subjects. Functional assays were performed to assess the duodenal epithelial digestive potential and barrier functionality.

**Results:** Organoid formation efficiency was similar between control-derived epithelial organoids (CDEOs) and MASH-derived epithelial organoids (MDEOs) (71% and 69%, respectively). Despite global heterogeneity in growth patterns, MDEOs frequently exhibited cystic spheroid morphology. MDEOs displayed altered digestive homeostasis associated with reduced mature absorptive cell fate, but they retained their lipid metabolic capacity, possibly mediated by lipid oxidation in stem/progenitor cells. Additionally, MDEOs misexpressed components of tight and adherens junctions and desmosomes compared to controls. However, MDEOs maintained pore and leak pathway integrity, indicating that the duodenal epithelial barrier remained functionally preserved under tested conditions.

**Conclusions:** This study provides evidence that the duodenal epithelium of MASH patients exhibits significant alterations in its digestive and barrier functions. This study sheds light on the intricate dynamics of duodenal epithelial alterations in MASH, highlighting potential therapeutic avenues for restoring intestinal homeostasis.

## INTRODUCTION

Metabolic dysfunction-associated steatotic liver disease (MASLD), formerly known as nonalcoholic fatty liver disease (NAFLD), is the leading cause of chronic liver diseases worldwide (1), while metabolic dysfunction-associated steatohepatitis (MASH), represents its advanced and progressive form (2–4).

Although MASH constitutes the hepatic manifestation of the metabolic syndrome, its development and progression involve multiple signaling pathways arising from different body systems (4), primarily, the adipose tissue and the gut. Furthermore, compelling evidence suggests that the gut-liver axis is intricately linked not only with the progression but also with MASH disease’s development. One of the key hallmarks is the disruption of the intestinal barrier integrity (5) leading to the passage of bacteria and their metabolic products into the portal system which ultimately worsen hepatic inflammation and drive metabolic alterations. More specifically, recent findings have shown that excessive fructose intake and high fat diet (saturated fat) promote intestinal barrier’s disruption (6), increase villus length expanding therefore nutrient absorption and adiposity (7, 8), and reduce mucous layer’s thickness (9), overall leading to systemic low-grade inflammation (10, 11). These findings emphasize the putative role of the small intestinal epithelium in the development and disease progression of MASH, but the fact that the above findings result from studies on mouse models is a limitation in fully understanding the mechanisms involved. As a result, the potential disruption of gut epithelium homeostasis in MASH patients is still incompletely studied, thereby its investigation could be of great interest in deciphering potential therapeutic targets at the level of intestinal mucosa. The breakthrough of intestinal organoid technology derived from individual patients has currently revolutionized translational research allowing the three-dimensional (3D) growth of tissue in cell culture derived from multipotent epithelial stem cells. The major advantage of 3D-organoids (unlike human primary epithelial cell models) resides in its capacity to self-organize and renew allowing to faithfully recapitulate the features and functions of the intestinal villous-crypt unit by maintaining cell diversity, barrier function, genetic and epigenetic pattern that are highly similar to in vivo (12) as already demonstrated in other gastrointestinal diseases such as inflammatory bowel disease (13–15). Consistently, few studies (16, 17) have already established liver organoids derived from MASH patients to further decipher the underlying molecular mechanisms at the level of the liver, however, so far, this model has not yet been used for studying gut epithelium.

Herein, we sought to further study the putative gut epithelium homeostasis disruption by generating duodenal organoids derived from MASH patients and healthy subjects. To this end, we analyzed patient-derived organoid transcriptomes from these 2 groups to evaluate if specific intestinal pathways were differentially modulated. Our data reveal that the duodenal epithelium of MASH patients exhibits significant alterations in its digestive and barrier functions, regardless of both luminal nutrient and microbial content and the surrounding subepithelial compartment.

## RESULTS

### Generation of a living duodenal organoid biobank from human MASH patients

To investigate potential contribution of the duodenal epithelium to metabolic dysregulation associated with MASH, we generated an organoid biobank from human duodenum biopsies obtained from 16 biopsy-proven MASH patients (Figure 1A). Eligible MASH patients were adults with NAFLD activity score (NAS) ≥4 integrating the following parameters: steatosis, ballooning, and inflammation. Main inclusion and exclusion criteria are reported in Materials and Methods section while the type 2 diabetes status and total cholesterol levels of included patients are reported in Figure S1A. Duodenum biopsies obtained from 14 adult outpatients who underwent routine esophagogastroduodenoscopy in the setting of epigastric pain and gastroesophageal reflux disease were used as controls. In line with the clinical metabolic status of MASH patients, body mass index (BMI) and age substantially differed between Control and MASH subjects. Despite over-representation of females over males in patients, such sex bias was present in both clinical groups [28.5% versus (vs) 25% males in Controls and MASH patients, respectively] (Figure 1A). Starting from an initial amount of 3 biopsies per patient collected in the second part of the duodenum (D2 post-papillary), we successfully generated 71% (10/14) Control-derived duodenal epithelial organoid (thereby referred as CDEOs) and 69% (11/16) MASH-derived duodenal epithelial organoid (thereby referred as MDEOs) lines. Organoids were amplified and stored as a biobank in the frame of 2 months following initial seeding (Figure 1B). Intra- and inter-individual heterogeneity were observed for grown elements within organoid lines. However, MDEOs more frequently adopted a cystic spheroid-like morphology as compared to CDEOs (Figure 1C). Of note, following freezing/thawing processes, 7/10 CDEO and 6/11 MDEO lines were efficiently replated beyond passage 5. Using the Phalloidin-FITC compound on organoid sections, we first assessed cell polarity. Both CDEOs and MDEOs were made of lumen-oriented apically polarized cells (Figure 1D). Scanning electron microscopy further confirmed the presence of crypt-like domains in CDEOs as compared to less frequently protruded MDEOs (Figure 1E). On the luminal side, microvilli were present in organoids of both types of subjects (Figure 1E, insets). Such phenotype was maintained over passages (Figure S1B).

**Figure 1.**
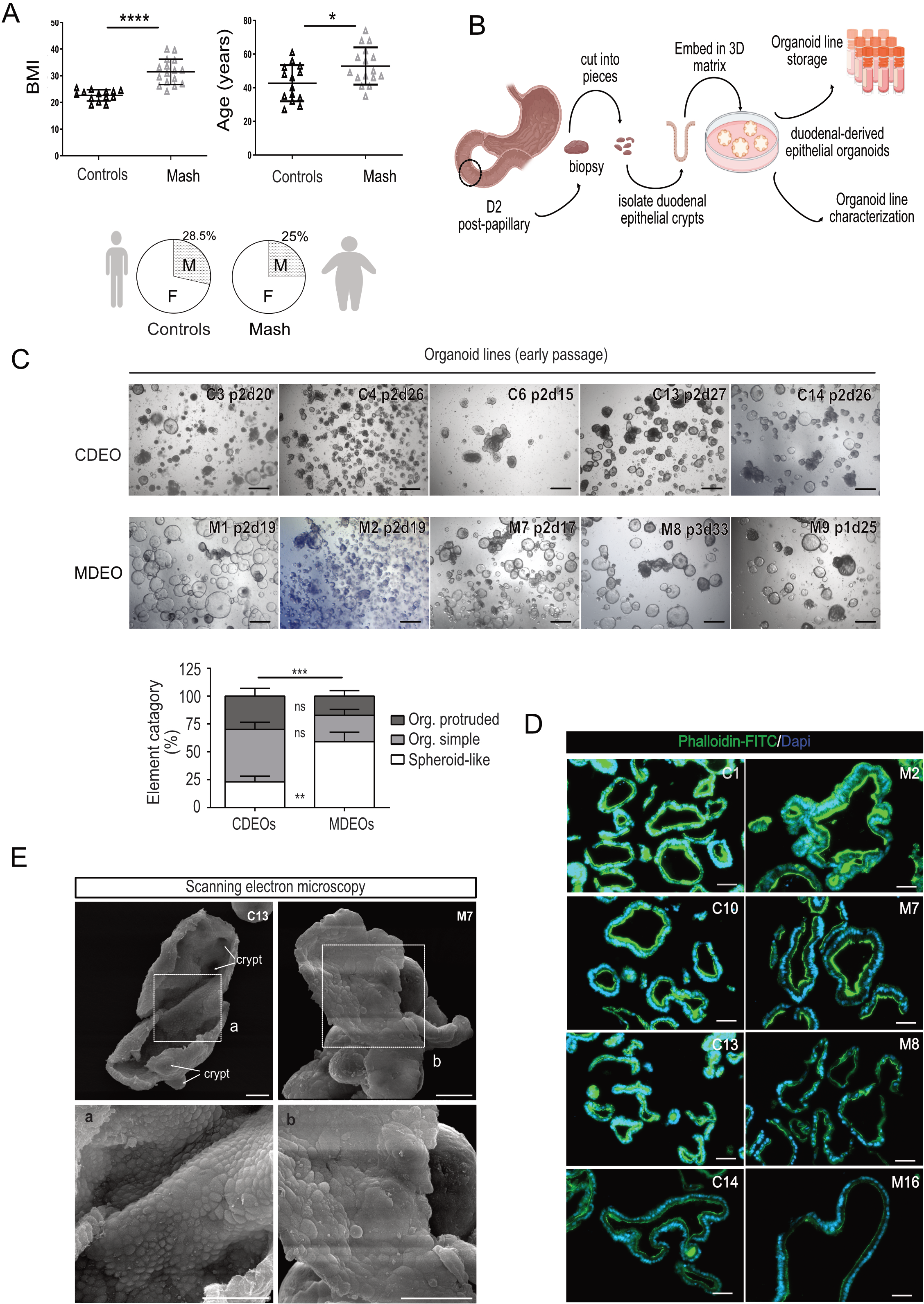
Generation of a living duodenal organoid biobank from human MASH patients. **A.** Demographic characteristics of patients selected for this study. Data are represented as means ± sd (standard deviation). Left panel: BMI (for body mass index): unpaired t-test: **** P< 0.0001; Right panel: age: unpaired t-test: *P< 0.05. Lower panel: Sex distribution (F: female; M: male). **B.** Scheme of the workflow from patient to organoid line storage (created with Biorender.com). **C.** Representative pictures of various CDEO and MDEO lines at the time of organoid line storage. Identity, passage, and day of culture are indicated for each organoid line. Scale bars: 500 µm. Lower panel: quantification of organoid complexity was determined for n=4 CDEO and 5 MDEO lines at the time of organoid line storage (1 to 3 passages). Org: organoid. Categories are expressed as the mean ± sem. Two-way ANOVA (interaction *** P< 0.001) followed by Sidak’s multiple comparisons test: MDEO vs CDEO spheroids: **P< 0.001; MDEO vs CDEO: organoid simple or organoid protruded: ns, not significant. **D.** Representative pictures showing apical polarity in CDEO and MDEO lines using Phalloidin-FITC. Nuclei were counterstained with DAPI. Scale bars: 50 µm. **E.** Representative picture of scanning electron microscopy showing crypt domains (arrows) and luminal side in CDEOs and MDEOs with insets (a, b). Scale bars: 50 µm.

Next, to determine whether MASH could be associated with any alterations in duodenal transcriptomic profiles, global transcriptomes of duodenal biopsies and duodenal-derived organoids isolated from Control and MASH patients (n= 12 and n= 13, respectively) were obtained by bulk RNA sequencing. First, we controlled tissue identity of biopsy and organoid samples by analyzing the expression of tissue-specific transcription factors. High levels of intestinal-specific CDX2 and CDX1 as well as PDX1 transcription factors were detected; in contrast to the foregut-related SOX2 factor, confirming the duodenal origin of samples (Figure S1C). We also excluded potential contamination of our organoid lines with CDX2-negative and CDH17-negative stem/progenitor cells from the submucosal Brunner’s glands (18, 19) (Figure S1D). For this purpose, immunofluorescence stainings were performed on patient biopsies and derived-organoid lines (Figure S1E, Figure S1F). Epithelial cells forming CDEOs and MDEOs were detected as CDH17-positive and expressed the intestinal crypt stem cell marker OLFM4 (Figure S1F). These data confirmed that organoid lines originated from duodenal crypts.

### MASH-derived organoids exhibit altered homeostasis

Next, we compared transcriptomic profiles of control and MASH-derived biopsies using the Degust software. However, no substantial differential gene expression could be observed between the groups when parameters were set to a False Discovery Rate (FDR) of 0.05 and absolute log2-fold change (FC) of 1. Then, we compared duodenal epithelium in both cohorts by analyzing the transcriptome of their derived organoids (n=7 CDEOs and 7 MDEOs) cultured under the same medium conditions for approximately 2 months after initial seeding. Using the same parameters as for biopsies’ analysis (i.e. FDR 0.05, log2-FC of 1), we identified 437 differentially expressed genes (DEGs), with 225 upregulated and 212 downregulated genes in MDEOs versus (vs) CDEOs (Figure 2A, 2B and 2C). Note that gene expression of Wnt signaling-associated intestinal stem cell markers (OLFM4, LGR5, AXIN2, SOX9, CD44) did not significantly differ between MDEOs vs CDEOs (Figure 2D) and cell proliferation was similar in both kinds of organoids (Figure 2E), indicating that stemness was preserved in the duodenal epithelium of MASH patients. However, analysis of the modulated “Hallmark” pathways revealed downregulation of the KRAS, hypoxia, inflammatory response, xenobiotic and lipid metabolisms, with concomitant upregulation of the estrogen response early/late and p53 pathways as well as the reactive oxygen species pathway (including the G6PD, PRDX4, AKR1B1, ARMCX1, LCN2 genes) in MDEOs vs CDEOs (Figure 2F, Figure S2A). Investigation of the associated “Biological process” pathways revealed deregulation of tissue homeostasis (59 modulated genes from the “homeostatic process” dataset list). Transcriptome analysis indicated downregulation of “phosphorylation” processes (p value 1.05e-12) in MDEOs (Figure 2G). This was associated with increased expression levels of several receptor-type tyrosine-protein phosphatases (PTPRU, PTPRM and PTPRS) reported to regulate the HIPPO/YAP and ERK signaling cascades (20, 21) (Figure 2G, Figure S2B). Altogether, these *in silico* studies suggested dysregulated signal transduction in the duodenal epithelium of MASH patients as compared to healthy subjects.

**Figure 2.**
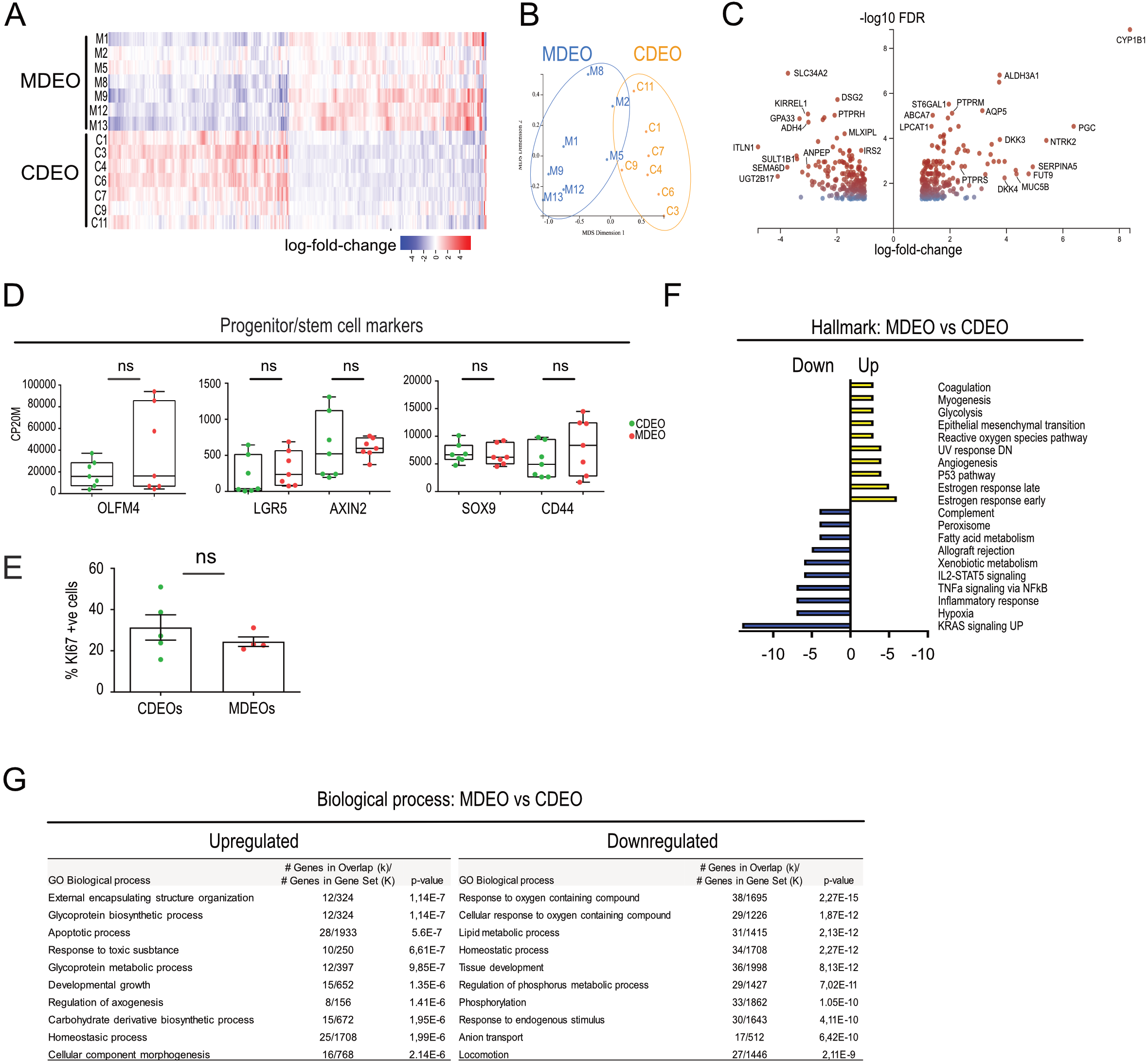
Characterization of tissue identity in patient biopsies and organoid lines. **A.** Heatmap of differentially expressed genes in CDEOs and MDEOs (log 2-fold change). **B.** Principal component analysis (PCA) plot of control (C) and MASH (M)-derived organoid transcriptomes. Each dot refers to an individual subject. **C.** Volcano plot showing transcriptome analysis of MDEOs vs CDEOs. Axes: x, log2-fold change; y, log10 of FDR (false discovery rate). Some genes involved in metabolism, cell signaling, and cell adhesion are highlighted. **D.** Expression levels of genes coding for progenitor/stem cell markers. CP20M: counts per kilobase of transcript per 20 million mapped reads. Each symbol corresponds to the value of an organoid line from an individual subject. Mann-Whitney test: ns, not significant. **E.** Cell proliferation index determined by the percentage of KI67-positive cells in organoids. Unpaired t test: ns, not significant. Each symbol corresponds to the value of an organoid line from an individual subject. **F.** GSEA-Hall mark for upregulated and downregulated gene lists in MDEOs vs CDEOs. p value is indicated. **G.** GSEA-Biological processes for upregulated and downregulated gene lists in MDEOs vs CDEOs.

### Modified digestive potential in duodenal-derived organoids of MASH patients

Strong downregulation of “response to oxygen containing compound” processes (p value 2.27e-15) was detected in MDEOs vs CDEOs (Figure 2G). Expression levels of genes associated with “lipid metabolic processes” (FABP2, DGKA, ACSL5, MOGAT3, SMPD3, GPA33 etc…) were reduced in MDEOs (Figure 3A). Several genes involved in carbohydrate metabolism were also downregulated, such as SGK1 regulating intestinal glucose absorption, the key regulator of lipid/carbohydrate’s metabolisms MLXIPL/CHREBP gene, as well as the pivotal Insulin-associated signaling molecule IRS2 (Figure 2C, S2A). Moreover, glucose transport mediated by solute carrier protein coding genes (SLCs) was altered, with downregulation of SLC2A2/GLUT2 and SLC5A1/GLUT1 and upregulation of the SLC2A10/GLUT10 (Figure S3A). In addition, expression of SLCs involved in amino acid, nucleoside, ionic transport was also deregulated in MDEOs (Figure S3A). Analysis of DEGs using the C8 “cell type signature” gene set indicated downregulation of immature, mature and late enterocytes-associated markers in MDEOs vs CDEOs (Figure 3B). Coherent with these results, we found de-enrichment of the “Brush Border membrane” and “Actin filament bundle” gene datasets in MDEOs vs CDEOs (Figure 3C). Moreover, 8 out of 29 genes of the human absorptive mature lineage list established by Gomez-Martinez et al. (22) were identified within the list of 212 downregulated genes in MDEOs vs CDEOs (Figure 3D). Regarding secretory lineages, we noticed slight upregulation of gastric mucins (MUC5B and MUC6) with concomitant decrease in the goblet marker MUC2 in MASH-derived organoids and biopsies (Figure 3E, Figure S3B). Altogether, these data suggested either bias commitment of stem cells toward the secretory vs absorptive lineages or reduced absorptive maturation of committed progenitors in MDEOs as compared to CDEOs. Next, having found evidence for altered digestive functions, we next compared the lipid metabolic potential of MDEOs and CDEOs. Organoids were stimulated 7 days post-replating with a mixture of oleic acid (OA) and palmitic acid (PA), two major fatty acids of the diet, for a further 4 days in culture. Globally, treatment promoted morphological conversion of spheroids into organoids, although MDEO lines exhibited heterogeneity (Figure 3F, left panel). Expression of genes involved in fatty acid cell entry or stem/progenitor markers did not appear significantly modulated by OA/PA challenge (Figure 3F, right panel). However, lipid challenge reduced expression of genes associated with fatty acid and triglyceride synthesis in a dose-dependent manner; and higher expression levels of ACLY, ACACA and FASN were detected in MDEOs vs CDEOs. Conversely, expression of CPT1A and HMGCS2 genes, involved in fatty acid beta-oxidation and ketone bodies synthesis, respectively, was stimulated by increasing concentrations of the mixture. MDEOs exhibited higher fatty acid oxidation potential as compared to CDEOs (Figure 3F, right panel). Taken together, these results indicated that MDEOs maintain their lipid metabolic capacity despite suggested lineage commitment bias, potentially through lipid oxidation mediated by stem/progenitor cells.

**Figure 3.**
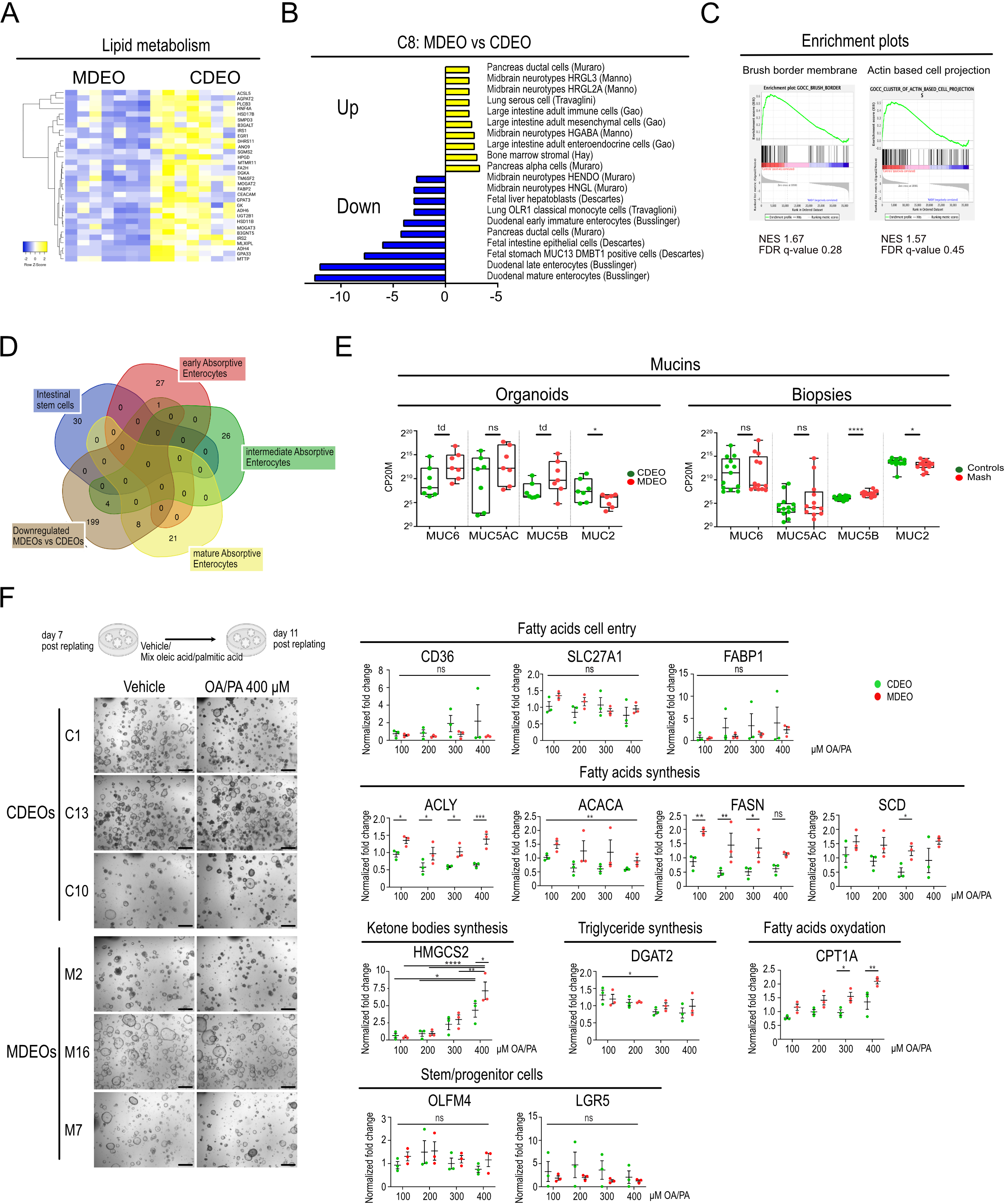
Transcriptome analysis of MASH-derived organoids reveals altered homeostasis. **A.** Heatmap showing differentially modulated genes involved in Lipid metabolism. **B.** GSEA-C8 “cell type signature” for upregulated and downregulated gene lists in MDEOs vs CDEOs. p value is indicated. **C.** GSEA showing de-enrichment of the “Brush border membrane” and “Actin based cell projection” dataset in MDEOs vs CDEOs. NES: Normalized Enrichment Score. FDR: False discovery rate. **D.** Venn diagram showing that genes expressed by mature absorptive enterocytes, as defined by Gomez-Martinez et al. (2022), are downregulated in MDEOs vs CDEOs. **E.** Expression levels of genes coding for various mucins. CP20M: counts per kilobase of transcript per 20 million mapped reads. Each symbol corresponds to the value of an organoid line or biopsy obtained from an individual subject. Mann-Whitney test: ****P < 0.0001; *P < 0.05; td: P<0.1; ns: not significant. **F.** Organoid challenge with mix oleic (OA) and palmitic acid (PA) for 4 days starting at 7 days post-replating. Left panel: Representative pictures of several organoid lines challenged with OA/PA (400 µM) or the vehicle. Scale bars: 500 µm. Right panel: Gene expression analysis by qRT-PCR of the indicated genes involved in lipid metabolism. Each symbol corresponds to a given organoid line. Two-way ANOVA tests with Sidak’s multiple comparisons. ****P < 0.0001; ***P < 0.001; **P < 0.01; *P < 0.05; ns: not significant.

### Misexpression of cell adhesion components in duodenal-derived organoids of MASH patients

Since mouse MASH model studies have reported disruption of intestinal epithelial barrier integrity (5), we analyzed expression levels of components of the apical junctional complexes. Expression of tight junction components (TJP1/ZO1, TJP2/ZO2 and JAM1) were found downregulated in MDEOs (Figure 4A, 4B, Figure S4A). Claudins, the other structural and functional components of tight junctions, also exhibited altered expression profiles. Nine out of 24 genes encoding members of this family of proteins were expressed at significant levels (CLDN1,2,3,4,7,12,15,18,23) in organoids. CLDN7, CLDN15, and to a lesser extent CLDN4, were downregulated in MDEOs (Figure 4A). Adherens junction components, which bring mechanical strength at cell-cell adhesion sites and contribute to cell polarization, were also dysregulated in MDEOs vs CDEOs. Indeed, cadherins (CDH1, CDH17, CDHR2) and catenins (CTNNB1, CTNNA1 and CTNND1), but not afadin or nectins, were significantly downregulated in MDEOs vs CDEOs (Figure 4A). Desmosome components (JUP, DSG2, DSC2, PKP2, PKP3) were also found downregulated in MDEOs as compared to CDEOs (Figure 4A). In line with transcriptomic data, the protein TJP1/ZO-1 was detected regularly spaced with punctuate apical staining in CDEOs but much less clearly visualized in MDEOs at any of the Z-stack focal planes analyzed (Figure 4C). Immunofluorescence studies using anti-DSC-2 antibodies, performed to visualize desmosomes at the intercellular junctions of the epithelium, suggested lower signal in MDEOs vs CDEOs (Figure S4B). Since loss of Desmocollin 2 in the mouse intestine results in desmosome ultrastructure alterations (23), we further analyzed the first most apical desmosome detected at the cell-cell junction by transmission electron microscopy (Figure 4D). This revealed shorter desmosomes in MDEOs as compared to CDEOs (Figure 4D). Altogether, these data indicated substantial intrinsic alterations of the cell-cell adhesion potential in MASH-derived organoids as compared to controls.

**Figure 4.**
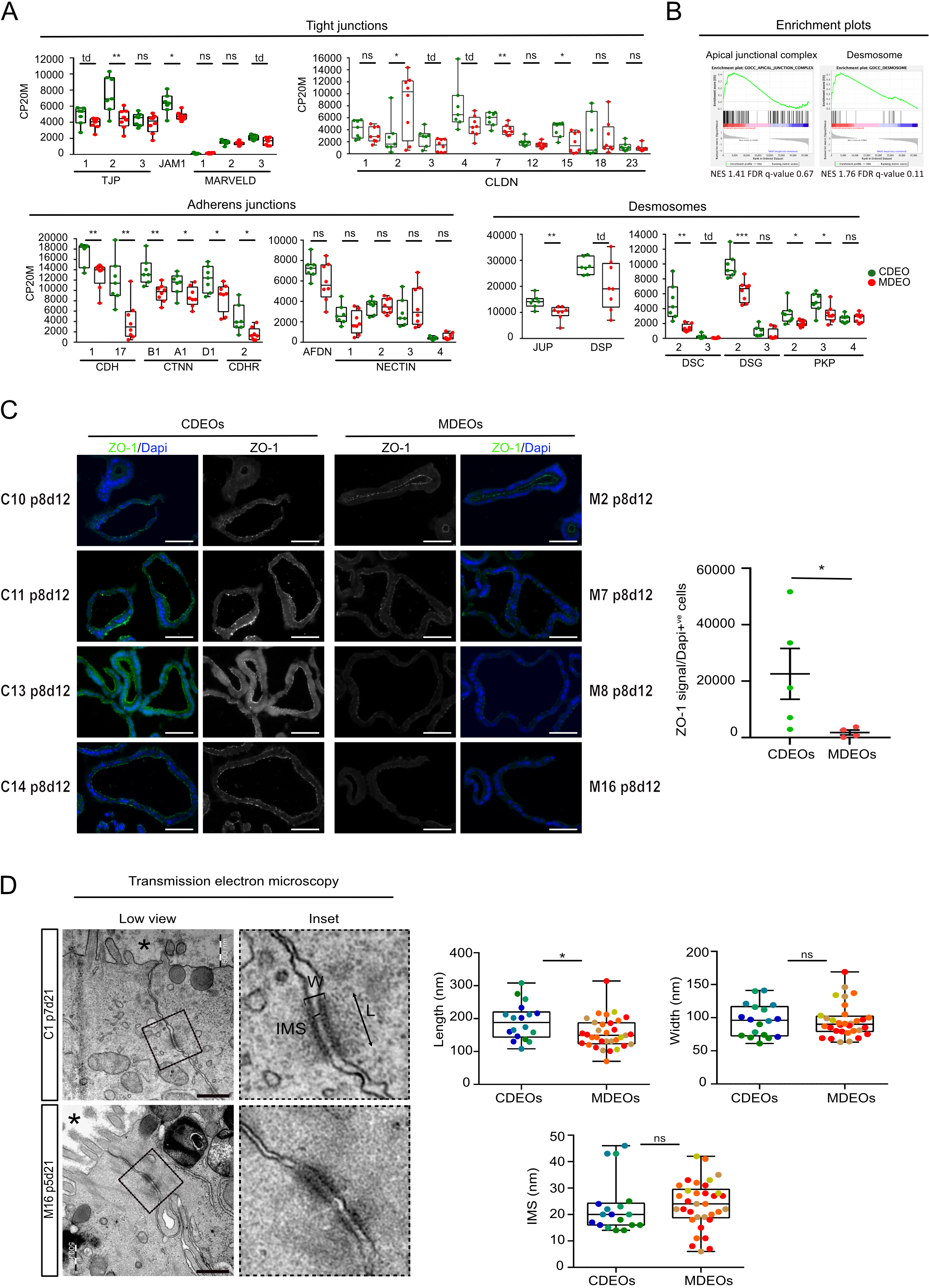
Transcriptome analysis of MASH-derived organoids reveals dysregulated expression of cell junction components. **A.** Expression levels of genes coding for components of tight and adherens junctions and desmosomes. CP20M: counts per kilobase of transcript per 20 million mapped reads. Each symbol corresponds to the value of an organoid line obtained from an individual patient. Mann-Whitney test: ***P < 0.001; **P < 0.01; *P < 0.05; td: P<0.1; ns: not significant. **B.** GSEA showing de-enrichment of the “Apical junctional complex” and “Desmosome” datasets in MDEOs vs CDEOs. NES: Normalized Enrichment Score. FDR: False discovery rate. **C.** Representative pictures of immunofluorescence showing discrete expression of ZO-1 in tight junctions in organoids. Nuclei counterstained with Dapi. Scale bars: 50 µm. Right panel: quantification of ZO-1 expression levels relative to the total number of cells. Each symbol refers to an individual organoid line (n=5 CDEOs, n=4 MDEOs). A mean of 267 cells were analyzed per organoid line. Mann-Whitney test: * P<0.05. **D.** Representative pictures of transmission electron microscopy showing altered desmosome architecture in MDEOs. Asterisks indicates the luminal side of organoids. Insets show higher magnification of desmosomes. W: width, L: length, IMS: intermembrane space. Scale bars: 500 nm. Quantification of the first most apical desmosome in MDEOs and CDEOs. Each symbol represents an individual desmosome and colors identify individual CDEO (n=4) or MDEO (n=4) lines. Unpaired t-test: * P<0.05, ns: not significant.

### The intestinal epithelial barrier is functionally preserved in MDEOs

To study the potential functional consequences of cell-cell adhesion modifications in MASH organoids, ultrastructure of the basolateral surface of organoids was analyzed by scanning electron microscopy. As shown in Figure 5A, meanwhile the basolateral surface was globally smooth for the various CDEOs, MDEOs exhibited higher density of crackles, this feature revealing looser cell-cell interactions in diseased-related organoids (Figure 5A). Moreover, transcriptome analysis revealed that the CLDN2 member, involved in tight junction formation, was upregulated in MDEOs vs CDEOs (Figure 4A). To confirm these data, we quantified its expression levels by RNAscope and found global increased levels in MDEOs, despite substantial heterogeneity among organoid lines (Figure 5B). Next, since CLDN2 is recognized as a mediator of leaky gut, we assessed epithelial barrier integrity in control and MASH-derived cultures (24). For this purpose, transepithelial electric resistance (TEER) of organoid lines was analyzed by culturing cells in 2D on transwells coated with Matrigel. As expected, establishment of a complete monolayer correlated with increased TEER over time in both kinds of cohorts (Figure 5C, left panel). Next, we investigated the tight and adherens junction reassembly potential following a calcium depletion challenge performed on fully grown monolayers. As expected, such treatment led to sharp reduction in TEER within 30 minutes whereas calcium replenishment was associated with TEER restoration within 2 hours (Figure 5C, right panel). However, these calcium challenges induced similar response regardless of the organoid line origin (Figure 5C, right panel). We also studied macromolecular paracellular permeability using the FITC-Dextran 4 kDa as a tracer molecule to explore the non-charged selective pathway. Cell permeability, calculated as the percentage of basolateral tracer detected after 24 hours, did not substantially differ between the two groups (Figure 5D). Together, these experiments indicated that despite modified cell adhesion properties of MDEOs vs CDEOs, permeability to ions, small molecules and macromolecules was globally preserved in MASH-derived epithelium under the tested culture conditions.

**Figure 5.**
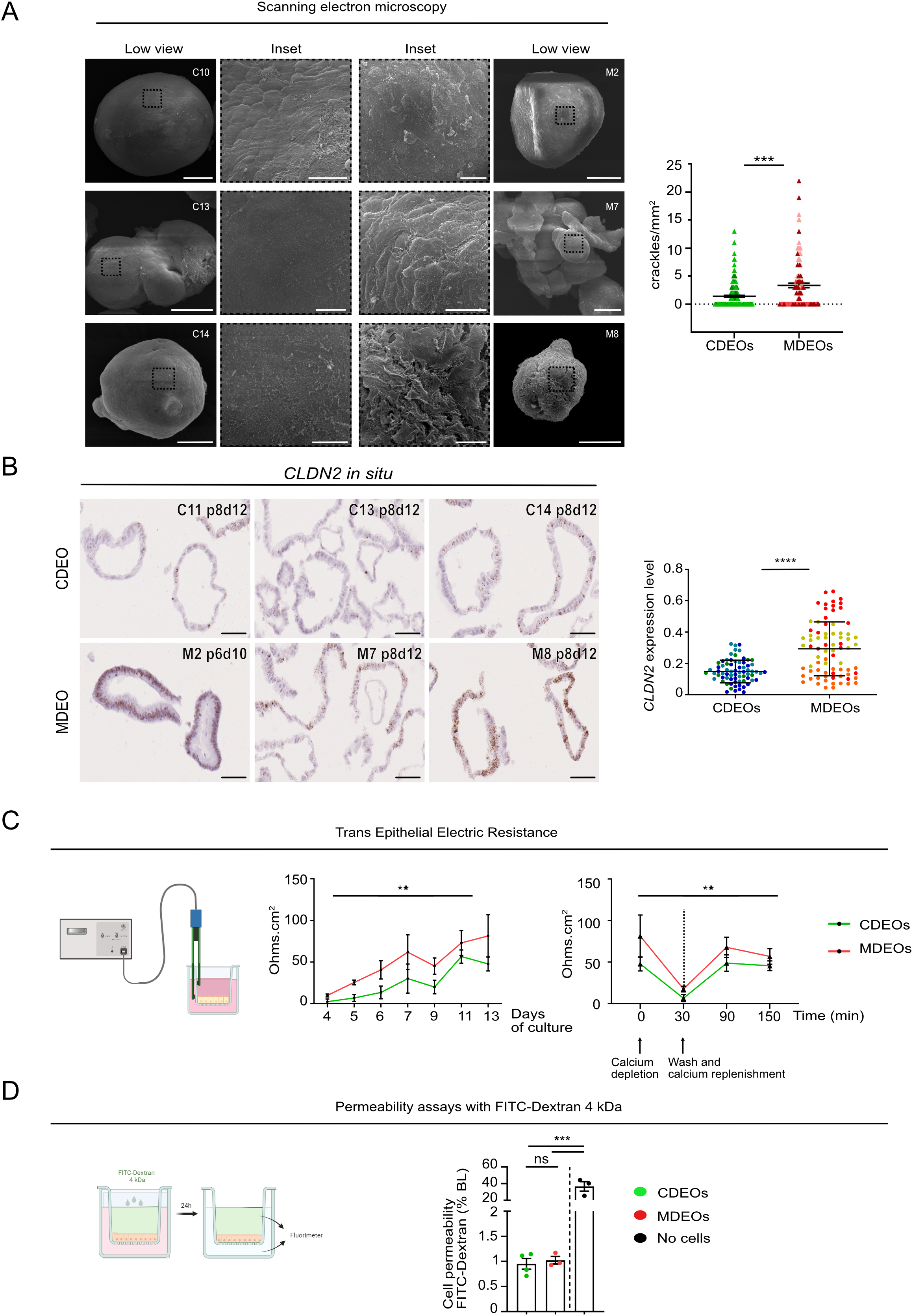
Preserved cell permeability of MASH duodenal-derived organoids. **A.** Representative pictures of scanning electron microscopy (inset: higher magnification) showing crackled areas at the basolateral surface of MDEOs. Scale bars: 50 µm, insets: 10 µm. Right panel: quantification of the number of crackles per mm2 of organoid surface performed on n=4 CDEOs and n= 3 MDEOs (35-45 pictures analyzed per organoid line). Mann-Whitney test: *** P < 0.001. **B.** Expression of the leaky gut associated CLDN2 gene detected in various CDEOs and MDEOs by RNAscope. Scale bars: 50 µm. Right panel: quantification of CLDN2 expression levels performed on n=3 CDEOs and n= 3 MDEOs (each line is identified with a different color) with a mean of 25 elements analyzed per organoid line. Data are expressed as the mean ± standard deviation. Mann-Whitney test: **** P<0.0001. **C.** Transepithelial electric resistance measurement performed on organoid lines grown on 2D cell inserts over time (left panel) or during a calcium depletion challenge (right panel) on n= 4 CDEOs and n=3 MDEOs. Left panel: Two-way ANOVA (time **** P< 0.0001, cohort ** P<0.01) followed by Tukey’s multiple comparisons test: MDEOs vs CDEOs d13: * P<0.05; CDEOs d4/d5 vs d11: ** P<0.01, d4/d5 vs d13: * P<0.05, d6 vs d13: * P<0.05; MDEOs d4 vs d7 : * P<0.05, d4 vs d11: ** P<0.01, d4 vs d13 *** P<0.001, d5 vs d11 * P<0.05, d5 vs d13 ** P<0.01. Right panel: Two-way ANOVA (time ** P< 0.01, cohort ** P<0.01) followed by Tukey’s multiple comparisons test: MDEOs vs CDEOs: ns, not significant; CDEOs at 30 min vs 90 min and CDEOs at 30 min vs 150 min: *P< 0.05. **D.** FITC-Dextran 4 kDa permeability assays performed on organoid lines grown on 2D cell inserts. Permeability is expressed as the percentage of tracer in the basolateral (BL) compartment after 24 hours. One-way anova test with tukey’s multiple comparisons. **** P<0.0001. CDEOs vs No cells: *** P< 0.001; MDEOs vs No cells: *** P< 0.001; MDEOs vs CDEOs: ns, not significant.

## DISCUSSION

In this study, we provide new evidence that the duodenal epithelium of MASH patients exhibits significant alterations in its digestive and barrier functions. This was observed in a culture model that isolated the epithelium to assess its intrinsic functional changes, irrespective of luminal nutrient and microbial contents, as well as the surrounding subepithelial compartment.

We used the organoid approach to generate a biobank of duodenal organoids obtained from control subjects and MASH patients. The success rate of this study, involving a limited number of individuals, was found to be similar in both cohorts, i.e., approximately 70%, indicating that MASH-derived stem cells had maintained their stemness capacity ex vivo. Consistent with previous reports on human liver organoid models for MASLD and MASH (16, 17, 25, 26), our study revealed both inter- and intra-organoid line heterogeneity in terms of morphology and gene expression within both types of organoids. Multiple differences in the origin of the duodenal samples obtained from individuals with distinct ages, clinical presentations, and dietary habits likely contribute to such variability. Nevertheless, despite this heterogeneity, a consistent phenotype was observed in MASH-derived organoids over passages. Of note, the fact that such differences were not detectable on biopsies at the transcriptomic level (except for the highly epithelial specific mucin-encoded genes) further highlights the potential of organoid technology for clinical studies. Persistence of the cystic morphology in MDEOs might be explained in part by differential expression of ionic and water flux solute carrier and aquaporin transporters, as revealed by bulk RNAseq.

Transcriptome profiling of MASH and Control-derived organoids revealed global dysregulation of tissue homeostasis associated with reduced response to organic compounds in disease-associated organoids, involving downregulation of lipid and xenobiotic metabolisms. These findings were suggestive of reduced absorptive cell lineage terminal differentiation, meanwhile secretory markers normally expressed in the anterior stomach (MUC5AC, MUC5B, MUC6, PGC), as well as several secretory or neuroendocrine markers (such as SCG5, PROX1, RETREG1, MAP1), were found upregulated in MASH organoids. Even though we failed to morphologically identify these neuroendocrine cells, our data are collectively suggestive of biased stem cell/progenitor lineage commitment in MASH organoids. Such observations are in line with recent single-cell RNAseq studies performed on mouse models showing the high degree of plasticity of the intestinal epithelium, able to modulate gene expression in response to acute or chronic nutrient challenges (27–29). Indeed, acute high-fat diet (HFD) leads to the appearance of new trajectories for secretory and enterocyte lineages associated with increase in the stem cell/progenitor pools (27). In a high-fat/high-sugar diet (HFHSD) model, scRNAseq studies have revealed a transcriptomic shift occurring in the small intestinal epithelium toward a more anterior cell type identity (28). In a human pathological context, increased expression of gastric mucins has been reported in the ulcer margins of ileal mucosa in Crohn’s disease (30). Accordingly, our data collectively suggest that MDEOs exhibit altered lineage differentiation compared to CDEOs when cultured under the same conditions. In the future, complementary scRNAseq studies should help to fully characterize the altered stem/progenitor trajectories in MASH-derived organoids.

MASH is associated with the accumulation of lipids in hepatocytes, leading to steatosis and progressive development of inflammation, which may result in the evolution of liver fibrosis (4). Previous ex vivo studies performed on human organoids generated from either induced pluripotent stem cells or direct liver bipotent ductal cells have reported impaired lipid oxidative metabolism leading to lipid droplet accumulation in MASH disease-related organoids or upon lipid challenge, as well as higher sensitivity to apoptosis following palmitic acid stimulation (16, 17, 25, 26, 31). In the present work, having provided transcriptomic evidence for dysregulated lipid metabolism in MASH duodenal epithelium, we explored their lipid metabolic potential upon free fatty acid stimulation. In line with mouse studies, fatty acids induced dose-dependent expression of CPT1A regulating fatty acid access to the mitochondrial matrix for β-oxidation and the HMGCS2 enzyme promoting mitochondrial ketone body formation and reported necessary to maintain intestinal stemness (29, 32, 33). However, we also found higher expression levels of genes involved in fatty acid biosynthesis in MASH vs control organoids. Overall, these experiments demonstrated the capacity of MASH organoids to metabolize fatty acids in response to a challenge, despite an observed global reduction in lipid metabolic processes. Future experiments meant to visualize the accumulation of lipid droplets in duodenal organoids would help to determine whether diseased organoids do exhibit preponderant lipid anabolic rather than catabolic processes as compared to controls.

Mouse models of MASH have provided evidence that liver inflammation is favored by gut-derived bacteria and bacterial metabolites following the initial disruption of intestinal barrier integrity (5). In humans, previous studies have shown less clear conclusions, but a recent meta-analysis review has correlated increased intestinal permeability in MASLD patients with the degree of liver steatosis (34). Moreover, a lifestyle that promoted weight loss successfully reversed the increased intestinal permeability and the extent of liver steatosis in obese patients (35). Various interdependent players, including the epithelial barrier itself, microbiota, immune, stromal, and enteric neurons, contribute to maintaining intestinal barrier integrity (36). Interestingly, the present study, focused on the sole epithelial player, has uncovered dysregulated expression of important components of cell-cell interaction, including the tight junction component CLDN2, one of the few members of this family of proteins, with pore-forming cation-selective channel properties. Of relevance, expression of this claudin, described as a leaky gut marker, is reported to be upregulated in several pathophysiological conditions such as inflammatory bowel diseases, coeliac disease, or irritable bowel disease (24). Nevertheless, it has recently been proposed that this claudin, detected in regenerative crypts of human inflammatory bowel diseases, would promote mucosal healing in a mouse experimental colitis model (37). Thus, the upregulation of CLDN2 in MASH vs control organoids observed in the present study might reflect a response to endogenous cellular stress occurring under the defined cultured conditions ; this would be coherent with the observed concomitant upregulation of a redox response in diseased organoids. However, another not mutually exclusive interpretation for CLDN2 upregulation in MASH organoids is also plausible. Indeed, differential expression of claudins has been reported along the crypt-villus axis; meanwhile Claudin 7 is highly expressed in differentiated cells of the villi, highest levels of Claudin 2 are detected in crypt domains (38). Therefore, the increased expression of CLDN2 in MASH organoids could be explained by relative amplification of the stem/progenitor pool as compared to control organoids. Certainly, additional studies are needed to fully decipher the function of this tight junction component in metabolic- and inflammation-related pathophysiological conditions. In addition to tight junction components, we also observed desmosome alterations and brittle basolateral cellular sides in MASH organoids by electron microscopy, suggesting reduced epithelial barrier potential. However, similar TEER and cell permeability to macromolecules were measured in MASH and control epithelia. In agreement with these findings, no significant differences were reported between severely obese and control subjects in intestinal permeability, despite evidence of morphological altered tight junctions in patient tissues, unless a lipid challenge was performed (39). Therefore, it is likely that the intrinsic transcriptomic and morphological alterations related to cell-cell adhesion detected in MDEOs vs CDEOs are not by themselves the unique drivers of the suggested in vivo gut permeability in MASH patients, but they certainly contribute to initiate the process. In addition, it is expected that the observed reduced production of secreted and glycocalyx-associated membrane-bound mucins can also further dampen epithelial barrier integrity and potentially modify the microbiota. Finally, the subtle default of the basolateral epithelial barrier observed by scanning electron microscopy could also favor access of stromal and immune cells to luminal content and thereby stimulate local inflammation. Further studies combining co-cultures of organoids with subepithelial cell types should help in the future to test this hypothesis.

In summary, despite the small number of samples involved in this study aimed at investigating the intrinsic characteristics of duodenal epithelia from MASH patients, we have revealed unique persistent alterations of disease-associated organoids. Since one of the most efficient treatments for MASH is lifestyle intervention, we anticipate that the duodenal organoid model could help test new therapeutic agents in the future to fully restore intestinal homeostasis.

## MATERIALS AND METHODS

### Human duodenal tissues

Human small intestine (duodenum) samples were obtained from biopsy-proven MASH patients who were included in a pilot study assessing the effect of an endoscopic duodenal mucosal resurfacing on liver histology outcome (40). Inclusion and exclusion criteria were the followings. Eligible patients were adults (aged ≥ 18 years) with histological evidence of definite steatohepatitis (per central expert pathologist reading of a liver biopsy obtained ≤ 6 months prior to the procedure) with an MAFLD activity score (NAS) of at least 4, including at least one point each for steatosis, lobular inflammation, hepatocellular ballooning; and fibrosis stage, per Brunt stage, of F0 to F3. Main exclusion criteria were as follows: evidence of another cause of liver disease; history of sustained alcohol ingestion defined as daily alcohol consumption > 30 g/day for males and > 20 g/day for females; previous gastrointestinal surgery such as Billroth 2, Roux-en-Y gastric bypass, or other similar procedures or conditions; for subjects with type 2 diabetes, no current use of insulin or GLP-1 analogs; and for type 1 diabetes, probable insulin production failure defined as fasting C-peptide serum < 1 ng/mL. We also obtained duodenum samples from outpatients who underwent routine esophagogastroduodenoscopy in the setting of epigastric pain and gastroesophageal reflux disease. Eligible patients were adults (aged ≥ 18 years) with body mass index < 25 with the following exclusion criteria: patients with type 2 diabetes, use of probiotics or antibiotics in the past 3 months. The study protocol was approved by the Medical Ethics Committee of CUB Hôpital Erasme, Hôpital Universitaire de Bruxelles (HUB); Université Libre de Bruxelles, Brussels, Belgium (approval number: B4062022000014). The study was conducted in accordance with ICH Good Clinical Practice Guidelines and the Declaration of Helsinki. All patients provided written informed consent. Clinical data for MASH patients are reported in Figure S1A.

### Crypt isolation and human duodenal organoid culture

Crypt isolation and organoid culture from human duodenal samples (3 biopsies) were performed based on the reported protocol (15). Biopsies were collected in the chelating buffer (CB) [constituted by cold sterile phosphate-buffered saline (PBS) supplemented with 2% sorbitol, 1% sucrose, 1% bovine serum albumin, gentamicin, and amphotericin B] (see Table 1-reagents). Briefly, villi were removed using fine scissors and the rest of the biopsy was further minced into 1 mm3 pieces and dissociated with 2 mM EDTA for 30 minutes at + 4°C with gentle rocking (90 rpm) to release crypts. Then, samples were centrifuged at 150 g for 5 minutes and resuspended in 5 mL of CB. Mechanical dissociation was carried out to release crypts by performing 40 times up and down pipetting with 5 mL pipette coated with fetal bovine serum. This fraction was next passed through a 100 µm-cell strainer. Filtered crypts were pelleted at 150 g for 5 minutes and suspended in Matrigel Growth factor reduced, Phenol-red free, LDEV-free onto 12-well plates as 100µl droplets. After Matrigel polymerization, samples were supplied with 700 µl culture medium and incubated at 37°C in a 5% CO2 cell culture incubator (Binder C150). The culture medium consisted of Advanced Dulbecco’s Modified Eagle’s medium supplemented with 5 U/ml penicillin/streptomycin cocktail, 40 µg/ml gentamycin, 2.5 µg/ml amphotericin B, 10 mM HEPES, 2 mM GlutaMAX, 1 mM N-acetyl cysteine, N2 supplement 1x, B-27 supplement without vitamin A 1x, 1 µg/ml human R Spondin-1, 100 ng/ml human Noggin, 50 ng/ml human recombinant epidermal growth factor, 10 nM Gastrin, 500 nM A83-01, 10 µM SB202190 and 10 mM of Nicotinamide (all indicated at the final concentrations, see Table 1-reagents). Culture medium was supplemented with 50% Wnt3A-conditioned medium (cell line Wnt-3A, ATCC, CRL-2647) containing 10% Fetal Bovine Serum, 1% Penicillin/Streptomycin, 10 mM HEPES, 2 mM GlutaMAX, as described (41). At initial seeding, 3,6 µM CHIR99021 and 2,5 µM thiazovivin were added to the medium. Culture medium was changed every two to three days. Organoid cultures were passaged every 15 to 20 days at a ratio of 1:5 using enzymatic dissociation with TrypLE express. Upon replating, culture medium was supplemented with 10 µM Y-27632 for the initial 2 days. For free fatty acid challenges, organoids were incubated 7 days after replating for 4 days with a free fatty acid mixture (stock solutions: oleic acid: 300 mM, palmitic acid: 250 mM final; 200 mM BSA) in complete culture medium. Images were acquired with a Moticam Pro camera connected to Motic AE31 microscope. Quantification of organoid morphology was performed in single-blinded with pictures acquired during culture between passages 1 to 6 on n = 8 CDEO and 7 MDEO lines. A mean number of 135 elements were analyzed per organoid line (with a minimum of 36 elements analyzed for the CDEO 6 line).

**TABLE 1.**
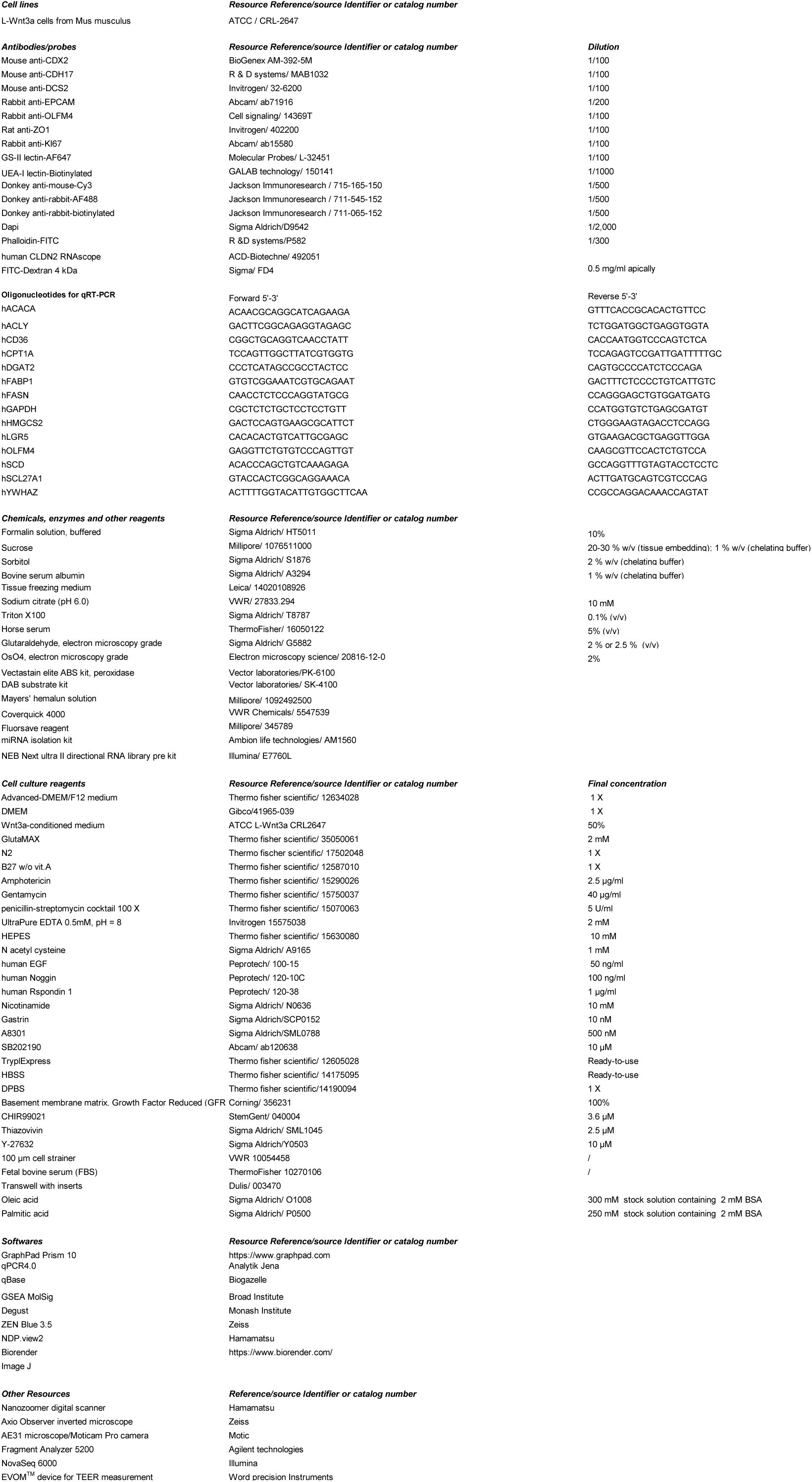
reagents.

### Tissue processing, immunohistochemical analysis and in situ hybridization

Biopsies were fixed in PFA 4% for 24 hours at +4°C and then processed for paraffin embedding (pathology laboratory, Institut Jules Bordet, Hôpital Universitaire de Bruxelles (HUB)). Samples were sectioned at 5 µm using the Sakura Tissue-tek AutoSection® microtome. Organoid culture samples were harvested, pre-fixed in 10% formalin solution during 30 minutes at room temperature (RT) and subsequently sedimented through 20% and 30% sucrose at 4°C before OCT embedding (Table 1-reagents). OCT samples were sectioned at 6 µm sections using a CM3050 cryostat (Leica Microsystems GmbH). For immunofluorescence or immunochemistry, antigen retrieval was performed by microwaving sections in 10 mM sodium citrate buffer, pH 6.0 before sample permeabilization (0.1 % Triton X-100 in DPBS). Nonspecific antibody binding was blocked for 1 hour at RT with blocking buffer (permeabilization buffer containing 5 % horse serum, 1 % BSA) before primary antibodies incubation overnight at + 4°C in blocking buffer. Sections were rinsed three times in PBS and incubated with secondary antibodies during 1 hour at RT. The ABC kits and substrate Kits (both from Vector Labs) were used for immunohistochemistry revelation. For fluorescence stainings, filamentous actin, lipids and nuclei were stained with phalloidin-FITC and DAPI, respectively. Slides were mounted with Fluorsave reagent. Antibodies and staining reagents are listed in Table 1-reagents. In situ hybridization experiments were performed on 6 µm-OCT embedded sections with the RNAscope kit (ACD-Biotechne) according to manufacturer instructions using the CLDN2 probe. Imaging was performed on a Zeiss Axio Imager M2 fluorescence microscope with a Zeiss Axiocam 503 mono camera and a Zeiss microscope Axio Observer 7 with axiocam MRN camera for immunofluorescence microscopy using Zen Pro software (Zeiss). Brightness, contrast, and picture size were adjusted using Zen lite software and fixed with a same setting for all pictures (Zeiss). Imaging for samples stained with antibodies or RNAscope probe by immunohistochemistry was performed with a Nanozoomer digital scanner (Hamamatsu). Quantification of CLDN2 expression levels was done with Q path 5.0 software by measuring the surface area (in µm2) of the probe signal over the surface area of the epithelial surface (in µm2) given by the hematoxylin signal. Results are presented as the ratio of CLDN2 signal/epithelium.

### Transmission electron microscopy (TEM) and scanning electron microscopy (SEM)

Organoids collected between day 16 to day 21 after replating (between passage 5 and 12) were washed with PBS and fixed with ice-cold glutaraldehyde 2% (Electron microscopy grade, Sigma #G5882) and post-fixed in 2% OsO4 in 0.1M cacodylate buffer (pH 7.2), serially dehydrated in increasing ethanol concentrations, embedded in Agar 100 resin (Agar Scientific Ltd, UK) and left to polymerize at 60°C for 2 days. Ultrathin sections (50-70 nm thick) were produced with a Leica EM UC6 ultra-microtome, collected on formvar-carbon-coated copper grids, and stained with uranyl acetate and lead citrate by standard procedures. Observations were made on a Tecnai 10 TEM (FEI) and images were captured with a Veleta CCD camera and processed with SIS iTEM (Olympus). Quantification of the dimensions of the most apical desmosome in organoids was performed using the NDPview2 software. For SEM, samples were fixed over night at 4°C in glutaraldehyde 2.5%, 0.1M cacodylate buffer (pH 7.2), and post-fixed in OsO4 (2%) in the same buffer. After serial dehydration, samples were dried at critical point and coated with platinum by standard procedures. Observations were made in a Tecnai FEG ESEM QUANTA 200 (FEI) and images processed by SIS iTEM (Olympus) software. Quantification of organoid crackles/mm2 was performed using the Image J software. Quantification was done on 4 CDEO and 3 MDEO lines, with 34-45 pictures being analyzed per organoid line.

### RNA extraction, RNA sequencing and Gene Set Enrichment Analysis (GSEA)

Organoid samples were collected at the indicated passage (p) and day (d): CDEO1 (p1d35), CDEO3 (p2d20), CDEO4 (p2d26), CDEO6 (p2d15), CDEO7 (p2d21), CDEO9 (p2d25), CDEO11 (p2d21), MDEO1 (p2d16), MDEO2 (p2d16), MDEO5 (p2d17), MDEO8 (p3d33), MDEO9 (p1d25), MDEO12 (p4d19), MDEO13 (p4d19). Total RNA from biopsies and organoids was extracted using the miRNA isolation kit (Ambion, Life Technologies, AM1560). RNA quality was checked using a Fragment analyzer 5200 (Agilent technologies). Indexed cDNA libraries were obtained using the NEBNext Ultra II directional RNA Library Prep Kit for Illumina® E7760L (NEW ENGLAND BioLabs Inc) following manufacturer recommendations. The multiplexed libraries were loaded onto a NovaSeq 6000 (Illumina) using an S2 flow cell and sequences were produced using a 200 Cycles Kit. Sequenced reads were trimmed for adaptor sequence (Trimmomatic-0.36). Paired-end reads were mapped against the human reference genome (GRCm38.p4/mm10) using the STAR_2.5.3a software to generate read alignments for each sample. After transcripts assembling, gene-level counts were obtained using HTSeq-0.9.1. Differentially expressed genes were identified with EdgeR method (Degust) and further analyzed using GSEA MolSig (Broad Institute) (42). Heatmaps were generated using Heatmapper (43).

### Gene expression analysis by qPCR

qRT-PCR was performed on total RNA extracted from organoid cultures using the miRNA isolation kit. A DNAse I treatment (Invitrogen) was used to remove potential contaminant DNA. cDNA was prepared using RnaseOUT and Superscript II according to the manufacturer’s protocol (Invitrogen). qPCRs were performed on the qTower 3 from Analytik Jena. Gene expression levels were normalized to that of reference genes (GAPDH, YWHAZ) and quantified using the qBase Software (CellCarta). Primer sequences are reported in the Table 1-reagents.

### Transepithelial electrical resistance measurements and paracellular permeability assay

To study cell permeability, organoid-generating cells were grown onto transwell inserts precoated overnight with 1 % Matrigel (500 000 cells seeded/well of p24 well plates, diameter: 6.5 mm, pore: 0.4 µm). Basolateral and apical chambers were filled with 600 µl and 200 µl of complete culture medium, respectively. Medium was changed every other day. Transepithelial electrical resistance was followed overtime using an epithelial volt-ohmmeter (EVOM device with STX4 electrodes). Calcium depletion assays were performed on fully confluent cultures (reached between day 13-15) by adding 2 mM EGTA in HBSS buffer for 30 minutes. Then, after wash with DPBS, cells were cultured in complete medium (containing 1.05 mM calcium chloride) for a further 150 minutes. Read data were corrected by subtracting the value of the blank corresponding to Matrigel-precoated wells containing no organoids and finally expressed as ohm.cm^2^. Experiments were repeated at least twice for each organoid line, with 3 measures/well at any time point for each well. Paracellular permeability was determined by measuring the diffusion of FITC-Dextran 4 kDa, initially added apically at 0.5 mg/ml to the culture medium, on the basolateral pole after 24 hours. Fluorescence was detected with the Microwin software on Mithras LB940 reader (Berthold Technologies) at 528 nm. Wells precoated with 1 % matrigel without cells were used as controls. Experiments were repeated at least twice for each organoid line, each measure was performed in triplicate.

### Statistical analysis

Statistical analyses were performed with Graph Pad Prism 10. All experimental data are expressed as mean ± standard error of the mean (sem) unless indicated in Figure legends. The significance of differences between groups was determined by appropriate parametric or non-parametric tests as described in Figure legends.

## Data availability

The datasets generated and analyzed during the current study are available in the GEODATASET repository [GEO Accession GSE 268518].

All authors had access to the study data and had reviewed and approved the final manuscript.

## Abbreviations

MASH: Metabolic dysfunction-associated steatohepatitis
CDEOs: control-derived epithelial organoids
MDEOs: MASH-derived epithelial organoids

## ACKNOWLEDGMENTS

We thank Eric Quertinmont and Vjola Tafçiu at the Laboratory of Experimental Gastroenterology for their help with the collection and procession of samples and RNA extraction.

## AUTHOR CONTRIBUTION STATEMENT

AH, ML: study concept and design, acquisition of data, analysis and interpretation of data, statistical analysis, drafting of the manuscript (ms).

AL, FL, GD, MM, MV, DM, DPM: acquisition of data, analysis and interpretation of data, statistical analysis.

AH, CL, CM, JD, ET: performed patient recruitment, study concept and design, study supervision, critical revision of the ms, obtained funding.

MIG: study concept and design, acquisition of data, analysis and interpretation of data, drafting of the ms, study supervision, obtained funding.

## Supplementary figure legends

**Figure S1.**
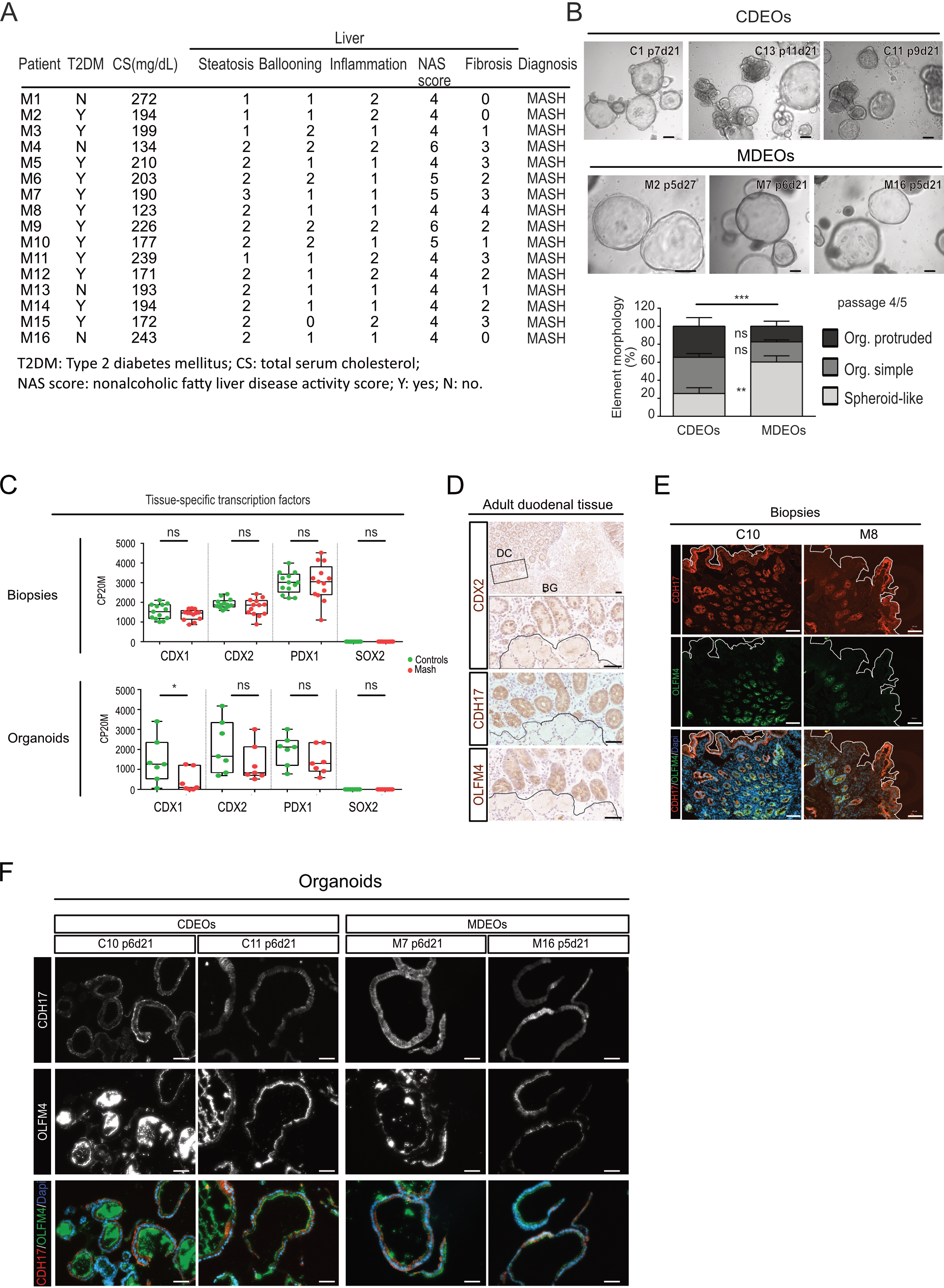
Generation of a living duodenal organoid biobank from human MASH patients. **A.** Table showing some biological parameters of MASH patients. **B.** Representative pictures of various CDEO and MDEO lines at advanced passage. Identity, passage, and day of culture are indicated for each organoid line. Scale bars: 150 µm. Righ panel: quantification of organoid complexity was determined for n=6 CDEO and 6 MDEO lines at passage 4-5. Org: organoid. Categories are expressed as the mean ± sem. Two-way ANOVA (interaction *** P< 0.001) followed by Sidak’s multiple comparisons test: MDEO vs CDEO spheroids: ** P< 0.001; MDEO vs CDEO organoid simple or organoid protruded: ns, not significant. **C.** Expression levels of genes coding for tissue-specific transcription factors. CP20M: counts per kilobase of transcript per 20 million mapped reads. Each symbol corresponds to the value of biopsy or organoid line obtained from an individual subject. Mann-Whitney test: * P< 0.01; ns, not significant. **D**. Immunohistochemistry showing expression of CDX2 and CDH17 in the duodenal epithelium and expression of the stem cell marker OLFM4 restricted to the duodenal crypts (DC). Brunner’s glands are delineated as BG. Scale bars: 50 µm. **E.** Immunofluorescence showing CDH17 and OLFM4 expression in biopsies. Nuclei counterstained with DAPI (merge image). The dotted line delineates the limit of the epithelium. Scale bars: 50 µm. **F.** Immunofluorescence showing CDH17 and OLFM4 expression in CDEO and MDEO lines. Identity and day of culture for each organoid line are indicated. Nuclei counterstained with DAPI (merge image). Scale bars: 50 µm.

**Figure S2.**
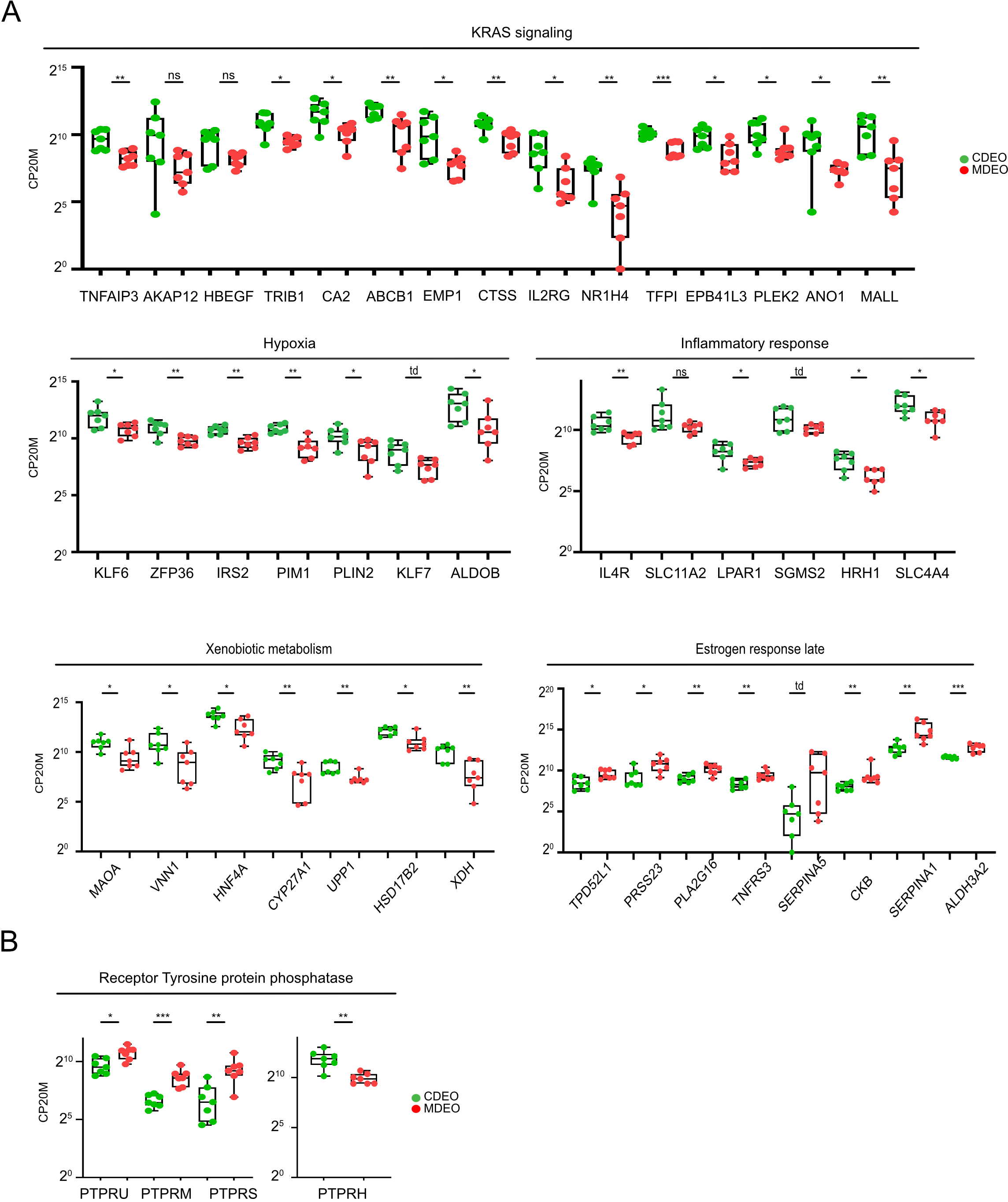
Altered cell signaling pathways in MDEOS vs CDEOs. **A.** Expression levels of genes involved in cell signaling processes. CP20M: counts per kilobase of transcript per 20 million mapped reads. Each symbol corresponds to the value of an organoid line obtained from an individual subject. Mann-Whitney test: *** P < 0.001; ** P < 0.01; * P < 0.05; td: P<0.1. **B**. Expression levels of genes coding for tyrosine protein phosphatases. CP20M: counts per kilobase of transcript per 20 million mapped reads. Each symbol corresponds to the value of an organoid line obtained from an individual subject. Mann-Whitney test: *** P < 0.001; ** P < 0.01; * P < 0.05.

**Figure S3.**
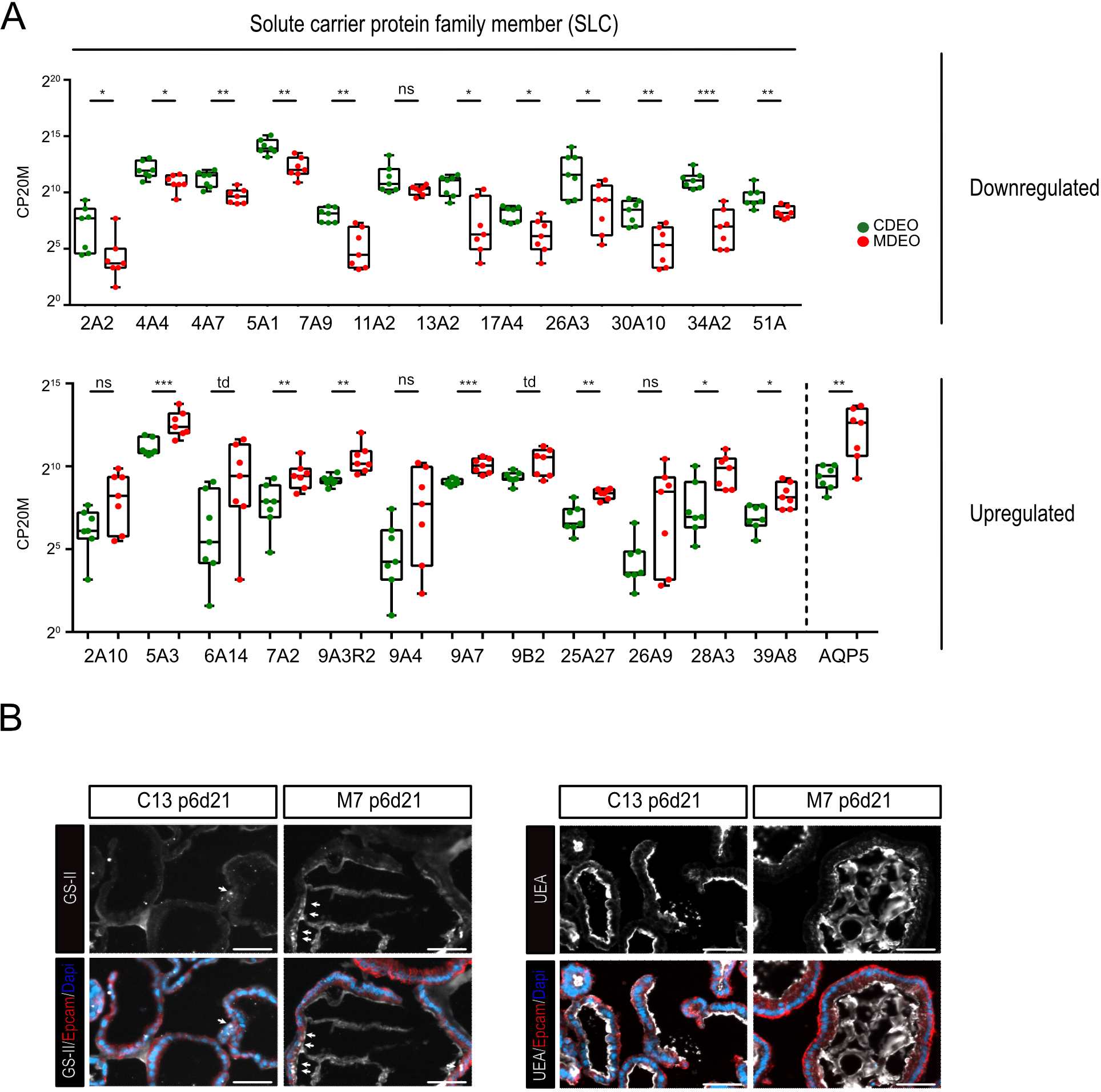
Transcriptome analysis of MASH-derived organoids reveals altered homeostasis. **A.** Expression levels of genes coding for differentially expressed solute carrier protein family members. CP20M: counts per kilobase of transcript per 20 million mapped reads. Each symbol corresponds to the value of an organoid line obtained from an individual subject. Mann-Whitney test: *** P < 0.001; ** P < 0.01; *P < 0.05; td (tendency): P<0.1; ns: not significant. **B.** Immunofluorescence showing altered expression of mucins in MDEOs vs CDEOs using GS-II and UEA-I lectins to label MUC6 and MUC2-expressing cells, respectively. Cell membranes shown with EPCAM and nuclei counterstained with DAPI. Arrows indicate cells labelled with GS-II. Scale bars: 50 μm.

**Figure S4.**
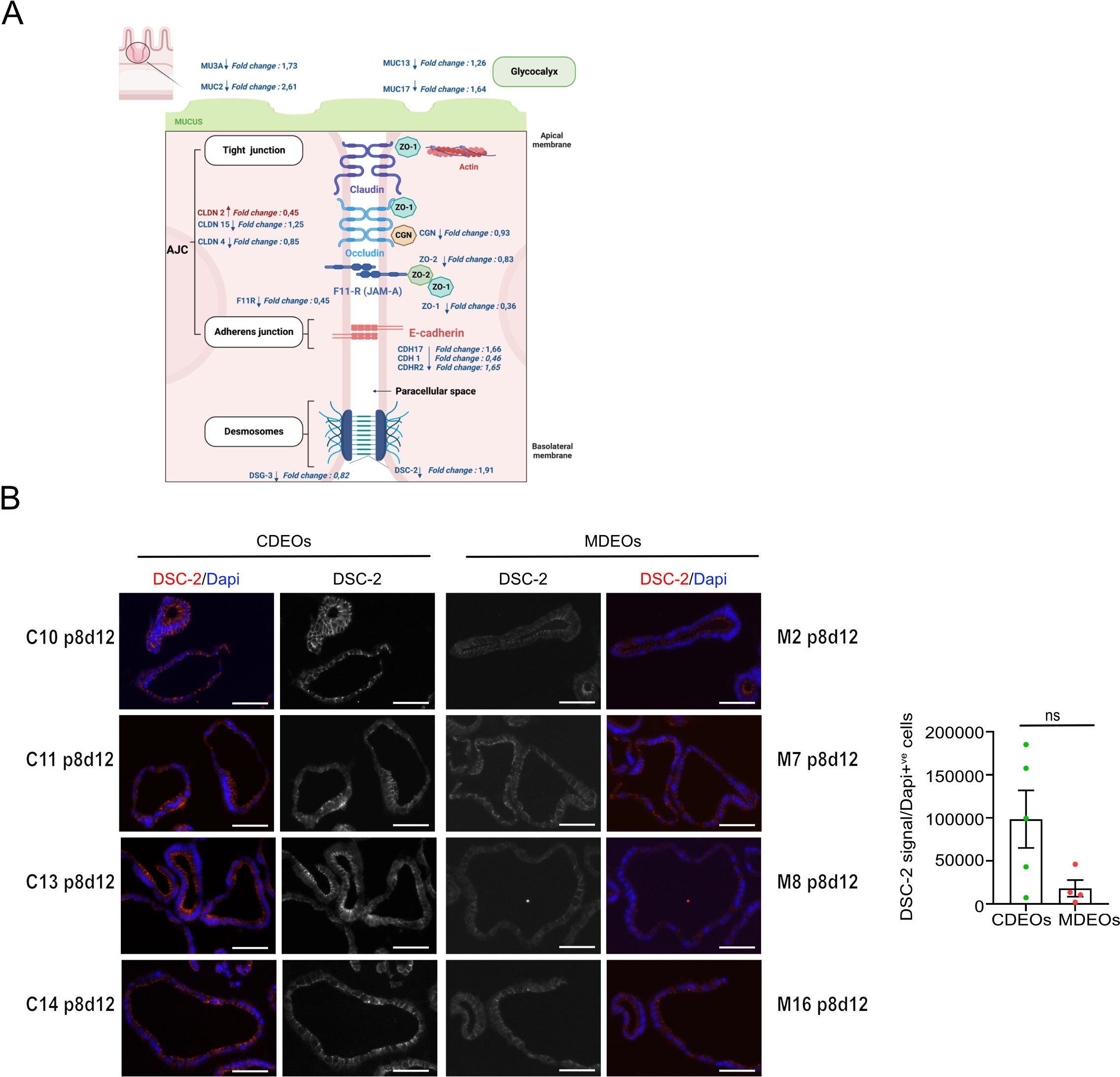
Transcriptome analysis of MASH-derived organoids reveals dysregulated expression of cell junction components. **A.** Schematic representation showing differentially modulated genes involved in cell adhesion. Log 2-fold change in MDEOs vs CDEOs is indicated. **B.** Representative pictures of immunofluorescence showing basolateral localization of DSC2 in organoids. Nuclei counterstained with Dapi. Scale bars: 50 µm. Right panel: quantification of DSC2 expression levels relative to the total number of cells. Each symbol refers to an individual organoid line (n=5 CDEOs, n=4 MDEOs). A mean of 267 cells were analyzed per organoid line. Mann-Whitney test: ns: not significant.

